## Supplementary information for "Stratification of viral shedding patterns in saliva of COVID-19 patients"

^1^interdisciplinary Biology Laboratory (iBLab), Division of Natural Science, Graduate School of Science, Nagoya University, Nagoya, Japan. ^2^School of Biomedical Convergence Engineering, Pusan National University, Yangsan, South Korea. ^3^Department of Science System Simulation, Pukyong National University, Busan, South Korea. ^4^Department of Mathematics, Pusan National University, Busan, South Korea. ^5^Lee Kong Chian School of Medicine, Nanyang Technological University, Singapore, Singapore. ^6^The Tokyo Foundation for Policy Research, Tokyo, Japan. ^7^International Research Center for Neurointelligence, The University of Tokyo Institutes for Advanced Study, The University of Tokyo, Tokyo, Japan. ^8^Department of Chemotherapy and Mycoses, National Institute of Infectious Diseases, Tokyo, Japan. ^9^Research Center for Drug and Vaccine Development, National Institute of Infectious Diseases, Tokyo, Japan. ^10^Department of Microbiology, University of Illinois at Urbana-Champaign, Urbana, IL, USA. ^11^Department of Statistics, University of Illinois at Urbana-Champaign, Urbana, IL, USA. ^12^Theoretical Biology and Biophysics, Los Alamos National Laboratory, Los Alamos, NM, USA. ^13^Institute of Mathematics for Industry, Kyushu University, Fukuoka, Japan. ^14^Institute for the Advanced Study of Human Biology (ASHBi), Kyoto University, Kyoto, Japan. ^15^Interdisciplinary Theoretical and Mathematical Sciences Program (iTHEMS), RIKEN, Saitama, Japan. ^16^NEXT-Ganken Program, Japanese Foundation for Cancer Research (JFCR), Tokyo, Japan. ^17^Science Groove Inc., Fukuoka, Japan. ^18^Division of Respirology, Rheumatology, Infectious Diseases, and Neurology, Department of Internal Medicine, Faculty of Medicine, University of Miyazaki, Miyazaki, Japan.

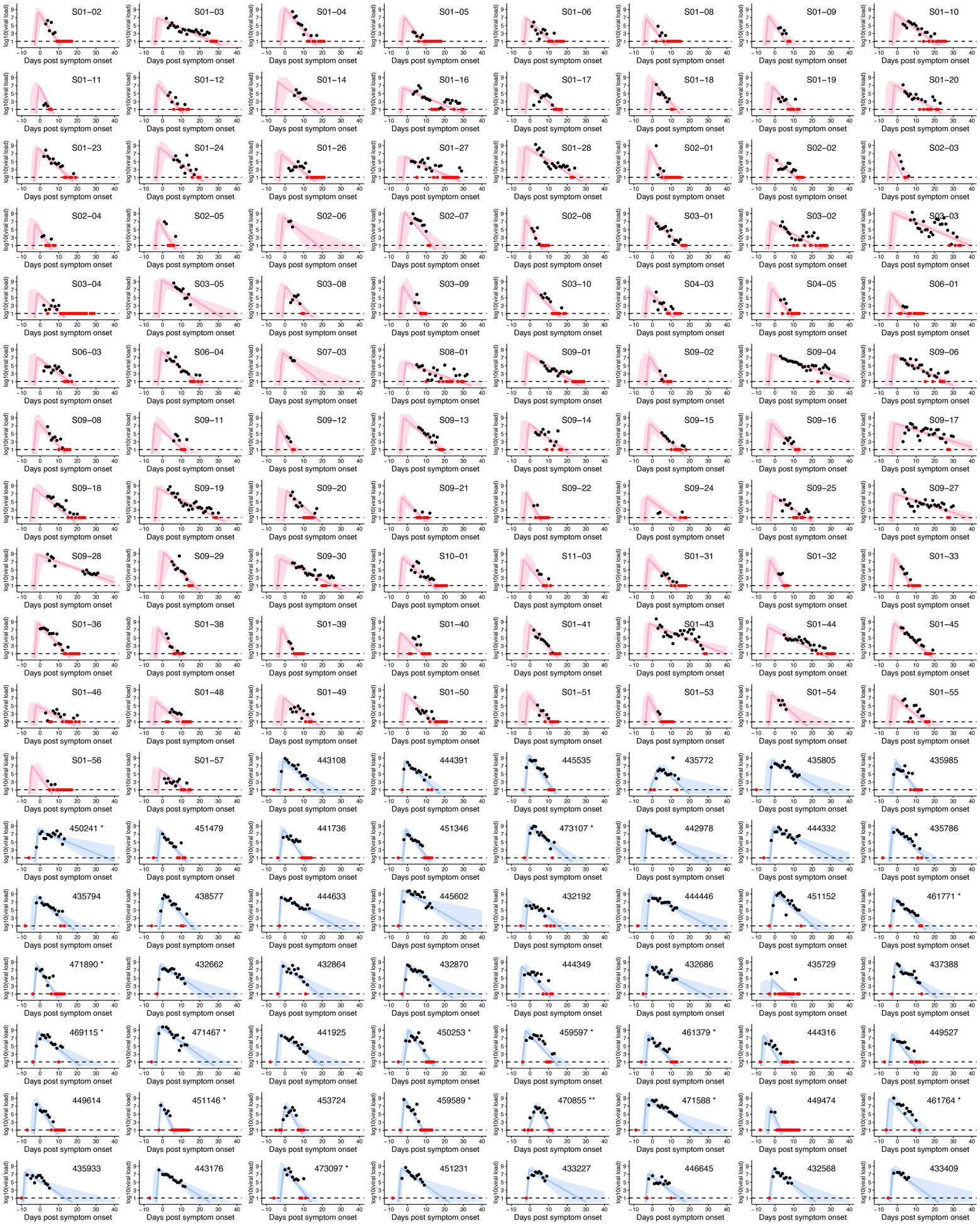

**Supplementary Figure 1.** **Reconstructed viral dynamics in saliva samples for individual participants:** The individual-level model fits to saliva RT-qPCR results, based on the target-cell-limited model described in Eqs.(1-2), are presented for the same cohorts shown in **Fig. 1**. Black and red closed dots represent measurements above and below the detection limit, respectively, with the black dashed line indicating the detection limit (1.08 log₁₀ copies/mL). Solid curves depict the reconstructed viral dynamics, while the shaded areas represent the corresponding 95% confidence intervals obtained using a bootstrap approach. Curves and shaded areas for individuals from the NFV clinical trial and the University of Illinois cohort are shown in pink and sky blue, respectively.

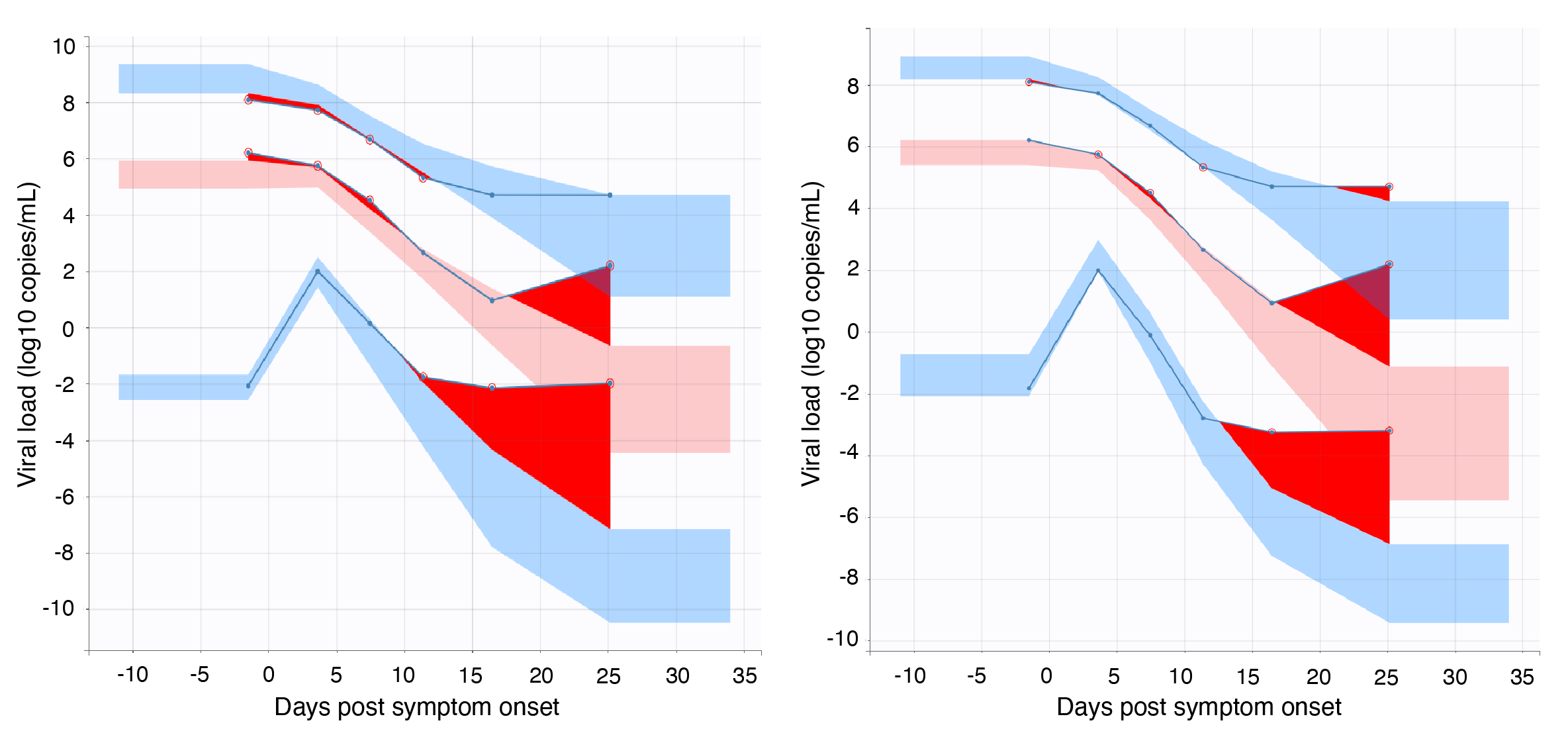

**Supplementary Figure 2. Visual predictive checks (VPC) for the viral load models:** Observed viral load data (red lines and points) are compared with simulated prediction intervals from the models (blue shaded areas, 95% prediction interval) for the target-cell limited model (left) and the immune effector cell model (right). The VPC shows that both models adequately capture the central tendency and variability of the observed data over time since symptom onset. In the later phase, some deviations are present, but these are similarly observed across both models.

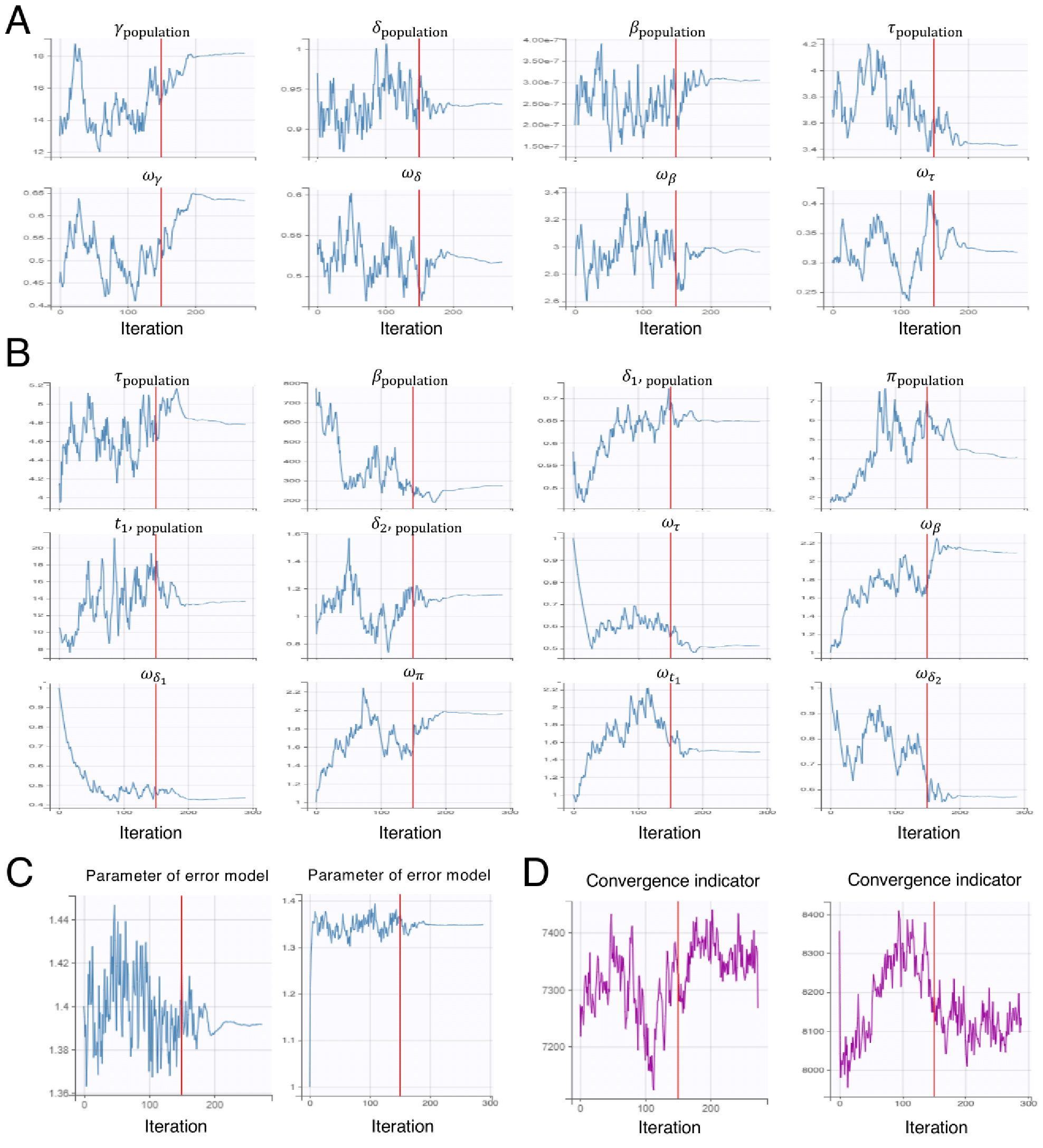

**Supplementary Figure 3. Convergence diagnostics of the SAEM algorithm: (A)(B)** Trajectories of population parameter estimates (with $\omega$ denoting the variance components of the random effects) over SAEM iterations are shown for the target-cell limited model **(A)** and the immune effector cell model **(B)**, respectively. In both models, parameter estimates stabilized after the burn-in period, indicating convergence. **(C)** Trajectories of the coefficient estimates in the linearized standard-error model are shown for the target-cell limited model (left) and the immune-effector cell model (right), respectively. **(D)** Convergence indicator values for the target-cell limited model (left) and the immune effector cell model (right) are shown, respectively. This indicator combines information on parameter stability and the Fisher information matrix. For both models, stabilization of the indicator after approximately N iterations indicates that the estimation procedure had reached convergence.

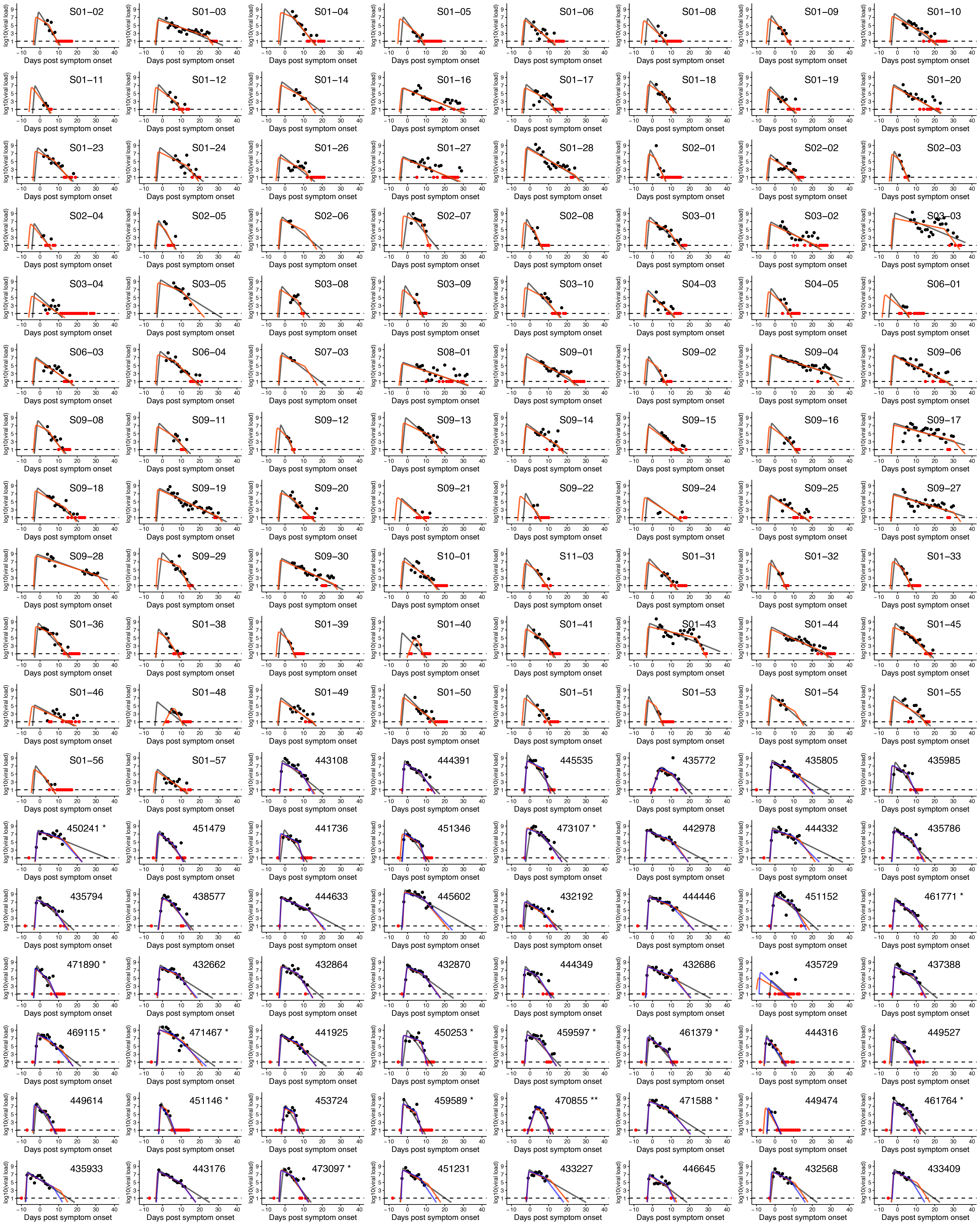

**Supplementary Figure 4. Comparison of three model fits to viral load in saliva samples for individual participants:** Three different individual-level model fits to saliva RT-qPCR results using the target-cell-limited model and the immune effector model described in Eqs.(1-2) and Eqs.(3-6), respectively, are shown. Black and red closed dots represent measurements above and below the detection limit, respectively, with the black dashed line indicating the detection limit (1.08 log₁₀ copies/mL). Black solid curves indicate the reconstructed viral dynamics using the target-cell-limited model presented in **Supplementary Fig 1**. The orange and blue curves indicate the estimated viral dynamics based on the immune effector model for all 144 individuals shown in **Fig 1** and for the 54 individuals only in the University of Illinois cohort, respectively. Namely, the blue dashed curves are the model fits presented in ^1^.

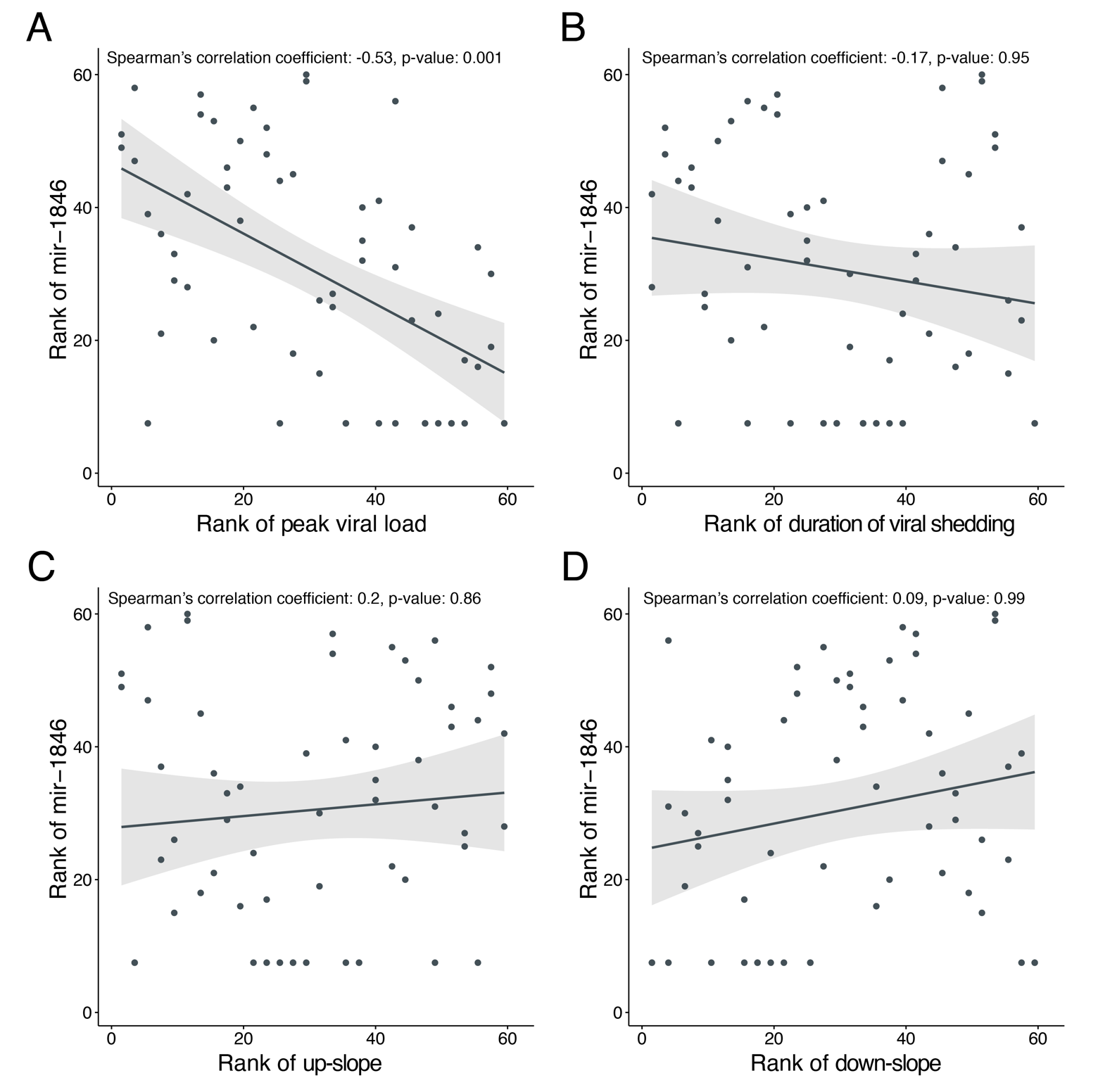

**Supplementary Figure 5.** **Correlation between mir-1846 level and the features of infection dynamics:** The correlation between the rank of mir-1846 and the rank of (A) the peak viral load, (B) the duration of viral RNA detection, (C) up-slope and (D) down-slope are shown, respectively. The Spearman’s correlation coefficients and p-values are presented in the top of each panel. The black solid line and the gray shaded area indicate results of the linear regression and their 95% confidence levels, respectively.

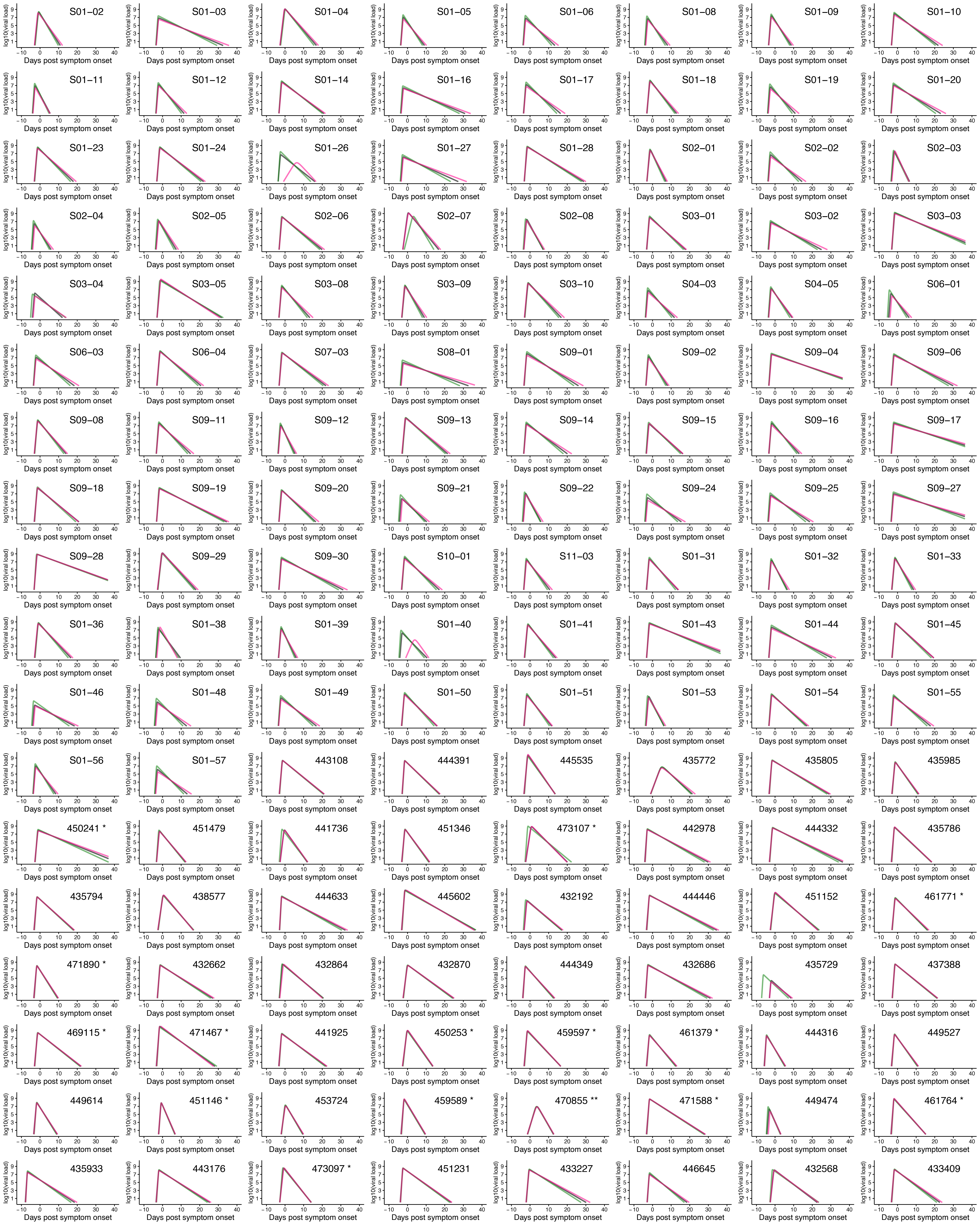

**Supplementary Figure 6. Sensitivity analysis of the detection limit for viral load data in the NFV cohort:** Three model fits to saliva RT-qPCR results using the target-cell-limited model described in Eqs.(1-2). The black, pink, and green curves correspond to the model fits obtained when the detection limit for the NFV cohort’s viral load data was set to 1.08 log_10_ copies/mL (same as the Illinois cohort), 2 log_10_ copies/mL, and 0 log_10_ copies/mL, respectively.

**Supplementary Table 1 | Daily symptom data for whole cohorts for each group**

| **Daily symptom** | **Group1 (N=46)** | **Group2 (N=61)** | **Group3 (N=37)** | **Overall (N=144)** |
| --- | --- | --- | --- | --- |
| Cough | 84.8% | 75.4% | 64.9% | 75.7% |
| Dyspnea | 32.6% | 24.6% | 37.8% | 30.6% |
| Nasal discharge | 58.7% | 67.2% | 59.5% | 62.5% |
| Sore throat | 63.0% | 65.6% | 59.5% | 63.2% |
| Nausea/vomiting/diarrhea | 52.2% | 68.9% | 54.1% | 59.7% |
| Myalgia | 39.1% | 57.4% | 45.9% | 48.6% |
| Olfaction abnormal | 41.3% | 52.5% | 59.5% | 50.7% |
| Fever | 45.7% | 24.6% | 32.4% | 33.3% |

**Supplementary Table 2 | Summary of parameter estimation by the model described in Eqs.(1-2)**

| **Parameters** | **Description** | **Unit** | $\boldsymbol{\vartheta}$**: Fixed effect (SE)*** | $\boldsymbol{\Omega}$**: SD of random effect (SE)*** |
| --- | --- | --- | --- | --- |
| $\beta$ | Rate constant for virus infection | (RNA copies/ml)^-1^day^-1^ | $2.08\times{10}^{-7}$ ($6.79\times{10}^{-8}$) | $2.79$ ($0.26$) |
| $\gamma$ | Maximum rate constant for viral replication | Day^-1^ | $13.2$ ($1.44$) | $0.45$ ($0.08$) |
| $\delta$ | Death rate of virus-producing cells | Day^-1^ | $0.97$ ($0.05$) | $0.54$ ($0.05$) |
| $\tau$ | Days from infection to symptom onset | Days | $3.73$ ($0.21$) | $0.31$ ($0.05$) |

* The parameter for patient $k$, $\vartheta_{i} \left( =\vartheta{\times e}^{\pi_{k}} \right)$ is represented as a product of $\vartheta$ (a fixed effect) and $e^{\pi_{k}}$ (a random effect). $\pi_{k}$ follows the normal distribution with mean 0 and standard deviation $\Omega$. SE: standard error.

**Supplementary Table 3 | Summary of parameter estimation by the model described in Eqs.(3-6)**

| **Parameters** | **Description** | **Unit** | $\boldsymbol{\vartheta}$**: Fixed effect (SE)*** | $\boldsymbol{\Omega}$**: SD of random effect (SE)*** |
| --- | --- | --- | --- | --- |
| $\beta$ | Rate constant for virus infection | (RNA copies/ml)^-1^day^-1^ | $7.74\times{10}^{-6}$ ($2.59\times{10}^{-6}$) | $2.04$ ($0.27$) |
| $\delta_{1}$ | Death rate of virus-producing cells | Day^-1^ | $0.58$ ($0.05$) | $0.52$ ($0.06$) |
| $\delta_{2}$ | Death rate of virus-producing cells by immune effector cells | Day^-1^ | $1.09$ ($0.13$) | $0.6$ ($0.11$) |
| $t_{1}$ | Days from symptom onset when immune effector cells start to work | Days | $10.5$ ($2.61$) | $1.44$ ($0.24$) |
| $\pi$ | Rate constant for virus production | Day^-1^ | $1.73$ ($0.59$) | $2.61$ ($0.31$) |
| $\tau$ | Days from infection to symptom onset | Days | $4.13$ ($0.28$) | $0.44$ ($0.06$) |

* The parameter for patient $k$, $\vartheta_{i} \left( =\vartheta{\times e}^{\pi_{k}} \right)$ is represented as a product of $\vartheta$ (a fixed effect) and $e^{\pi_{k}}$ (a random effect). $\pi_{k}$ follows the normal distribution with mean 0 and standard deviation $\Omega$. SE: standard error.

**Supplementary Table 4 | Features of reconstructed individual viral dynamics for each group**

| **Feature** | **Group1 (N=46)** | **Group2 (N=61)** | **Group3 (N=37)** | **Overall (N=144)** | **p-value** |
| --- | --- | --- | --- | --- | --- |
| Duration of virus shedding (days) | 11.5 (3.16)* | 17.4 (3.14) | 30.0 (5.8) | 18.8 (8.13) | <0.001 |
| Peak viral load (RNA copies/ml) | 6.94 (0.69) | 8.21 (0.56) | 7.95 (1.03) | 7.74 (0.93) | 0.020 |
| Up-slope (day^-1^) | 12.2 (1.65) | 11.2 (1.85) | 13 (1.18) | 12 (1.79) | <0.001 |
| Down-slope (day^-1^) | 1.53 (0.55) | 1.11 (0.24) | 0.57 (0.13) | 1.1 (0.51) | <0.001 |

* Mean (standard deviation)

**Supplementary Table 5 | Pearson’s correlation coefficients between clinical data and features of viral dynamics.**

| **Clinical data** | **Duration of viral RNA detection** | | **Peak viral load** | | **Up-slope** | | **Down-slope** | |
| --- | --- | --- | --- | --- | --- | --- | --- | --- |
|  | Pearson’s $r$ | p-value | Pearson’s $r$ | p-value | Pearson’s $r$ | p-value | Pearson’s $r$ | p-value |
| Age | -0.08 | 0.8359 | -0.16 | 0.5783 | 0.15 | 0.5880 | 0.08 | 0.8141 |
| Systolic blood pressure | 0.08 | 0.8359 | -0.04 | 0.9388 | 0.12 | 0.6448 | -0.16 | 0.4519 |
| Diastolic blood pressure | -0.06 | 0.8359 | -0.2 | 0.5152 | 0.14 | 0.5880 | -0.03 | 0.9204 |
| Pulse rate | -0.13 | 0.6307 | -0.05 | 0.9388 | 0 | 0.9710 | 0.01 | 0.9445 |
| SpO_2_ | -0.04 | 0.8359 | -0.06 | 0.9388 | 0.07 | 0.9178 | 0.01 | 0.9512 |
| Respiratory rate | 0 | 0.9770 | 0.02 | 0.9765 | 0.08 | 0.9178 | 0.04 | 0.8681 |
| White blood cell count | -0.07 | 0.8359 | 0 | 0.9868 | -0.01 | 0.9710 | 0.07 | 0.8610 |
| Neutrophil | -0.04 | 0.8359 | -0.02 | 0.9765 | 0.15 | 0.5880 | 0.05 | 0.8681 |
| Eosinophil | -0.04 | 0.8359 | -0.24 | 0.4083 | 0.16 | 0.5880 | -0.08 | 0.8386 |
| Basophil | 0.09 | 0.8359 | -0.05 | 0.9388 | -0.15 | 0.5880 | -0.14 | 0.4933 |
| Lymphocytes | 0.01 | 0.9770 | 0.04 | 0.9388 | -0.15 | 0.5880 | -0.02 | 0.9231 |
| Monocyte | 0.11 | 0.6993 | 0.08 | 0.8939 | -0.09 | 0.9166 | -0.06 | 0.8681 |
| Red blood cell count | 0.08 | 0.8359 | 0.07 | 0.8939 | 0.01 | 0.9710 | 0.01 | 0.9445 |
| Amount of hemoglobin | 0.26 | 0.1688 | 0.26 | 0.4083 | 0 | 0.9710 | -0.2 | 0.3194 |
| Hematocrit | 0.25 | 0.1688 | 0.18 | 0.5209 | 0.07 | 0.9178 | -0.2 | 0.3194 |
| Platelet count | -0.05 | 0.8359 | -0.13 | 0.6790 | 0.03 | 0.9710 | -0.04 | 0.8681 |
| CRP | -0.06 | 0.8359 | 0.01 | 0.9765 | 0.02 | 0.9710 | 0.13 | 0.4933 |
| Protein | 0 | 0.9770 | -0.1 | 0.8684 | 0.08 | 0.9178 | -0.03 | 0.9200 |
| Albumin | 0.22 | 0.1688 | 0.21 | 0.5152 | -0.15 | 0.5880 | -0.17 | 0.3756 |
| ALT | 0.24 | 0.1688 | 0.04 | 0.9388 | 0.14 | 0.5880 | -0.24 | 0.2570 |
| AST | 0.26 | 0.1688 | 0.16 | 0.5783 | 0.1 | 0.8732 | -0.23 | 0.2570 |
| γ-GTP | 0.16 | 0.4469 | -0.07 | 0.8939 | 0.18 | 0.5880 | -0.21 | 0.3194 |
| ALP | 0.21 | 0.1791 | -0.07 | 0.8939 | 0.26 | 0.4708 | -0.27 | 0.2570 |
| LDH | 0.12 | 0.6676 | -0.01 | 0.9765 | 0.06 | 0.9178 | -0.15 | 0.4661 |
| Bilirubin | 0.22 | 0.1688 | 0.15 | 0.5783 | 0.03 | 0.9710 | -0.19 | 0.3194 |
| CK | 0.08 | 0.8359 | 0.01 | 0.9765 | -0.04 | 0.9710 | -0.12 | 0.4971 |
| Na | -0.02 | 0.9374 | -0.12 | 0.7050 | 0.02 | 0.9710 | -0.05 | 0.8681 |
| K | -0.02 | 0.9374 | -0.08 | 0.8939 | 0.06 | 0.9178 | 0.07 | 0.8610 |
| Cl | -0.07 | 0.8359 | -0.19 | 0.5209 | 0.06 | 0.9178 | -0.04 | 0.8681 |
| BUN | 0.22 | 0.1688 | -0.03 | 0.9388 | 0.13 | 0.6448 | -0.25 | 0.2570 |
| CRE | 0.22 | 0.1688 | 0.18 | 0.5209 | 0.03 | 0.9710 | -0.14 | 0.4933 |
| Blood glucose | 0.04 | 0.8359 | -0.12 | 0.7050 | 0.15 | 0.5880 | -0.06 | 0.8681 |
| HbA1c | -0.04 | 0.8359 | -0.13 | 0.6790 | 0.07 | 0.9178 | 0.04 | 0.8681 |
| Procalcitonin | -0.14 | 0.5327 | 0.05 | 0.9388 | -0.18 | 0.5880 | 0.19 | 0.3194 |
| Fibrinogen | -0.08 | 0.8359 | 0.02 | 0.9765 | 0.04 | 0.9710 | 0.13 | 0.4933 |
| PT | 0.16 | 0.4469 | 0.03 | 0.9388 | -0.02 | 0.9710 | -0.12 | 0.4971 |
| APTT | -0.05 | 0.8359 | 0.08 | 0.8939 | 0.01 | 0.9710 | 0.13 | 0.4971 |
| PT-INR | -0.22 | 0.1688 | -0.01 | 0.9765 | -0.01 | 0.9710 | 0.18 | 0.3655 |
| D-dimer | -0.06 | 0.8359 | -0.13 | 0.6790 | 0.13 | 0.6448 | -0.01 | 0.9512 |

**Supplementary Table 6 | Salivary micro-RNAs obtained from 60 specimens of 30 participants**

| **Micro-RNA** | **Group1 (N=10)** | **Group2**  **(N=10)** | **Group3**  **(N=10)** | **Overall**  **(N=30)** |
| --- | --- | --- | --- | --- |
| mir-10 | 45.4 (28.98) | 27.5 (11.41) | 34.25 (23.71) | 35.72 (23.43) |
| mir-103 | 0.12 (0.27) | 0.42 (0.85) | 0.11 (0.23) | 0.22 (0.54) |
| mir-1180 | 2.16 (4.39) | 0.94 (1.3) | 0.81 (1.45) | 1.3 (2.79) |
| mir-1226 | 0.85 (1.3) | 0.63 (0.85) | 0.44 (0.74) | 0.64 (0.99) |
| mir-1253 | 0.04 (0.16) | 0 (0) | 0 (0) | 0.01 (0.09) |
| mir-1255 | 0.09 (0.23) | 0 (0) | 0 (0) | 0.03 (0.14) |
| mir-128 | 0 (0) | 0.03 (0.12) | 0.01 (0.05) | 0.01 (0.07) |
| mir-1296 | 1.25 (2.41) | 0.41 (0.63) | 0.46 (0.78) | 0.71 (1.53) |
| mir-130 | 1.02 (1.98) | 0.74 (0.91) | 1.04 (1.12) | 0.93 (1.4) |
| mir-145 | 0 (0) | 0.01 (0.06) | 0.02 (0.1) | 0.01 (0.07) |
| mir-146 | 9.72 (4.52) | 11.41 (6.49) | 10.98 (6.22) | 10.71 (5.76) |
| mir-148 | 28.7 (5.9) | 31.47 (6.88) | 29.53 (10.38) | 29.9 (7.91) |
| mir-154 | 0.6 (1.95) | 1 (1.68) | 0.82 (1.88) | 0.81 (1.82) |
| mir-16 | 174.2 (34.41) | 178.55 (46.78) | 184.2 (37.33) | 178.98 (39.39) |
| mir-17 | 0 (0) | 0.02 (0.11) | 0 (0) | 0.01 (0.06) |
| mir-181 | 0.93 (1.36) | 0.99 (1.1) | 1.28 (2.15) | 1.07 (1.58) |
| mir-185 | 7.39 (7.39) | 5.04 (2.31) | 7.87 (13.71) | 6.77 (9.02) |
| mir-19 | 14.13 (8.99) | 17.23 (12.16) | 11.49 (6.35) | 14.28 (9.6) |
| mir-191 | 2.87 (5.2) | 1.22 (1.54) | 1.51 (1.68) | 1.87 (3.3) |
| mir-192 | 0.14 (0.46) | 0.07 (0.3) | 0.04 (0.13) | 0.08 (0.32) |
| mir-193 | 0.16 (0.7) | 0 (0) | 0 (0) | 0.05 (0.4) |
| mir-204 | 0.21 (0.6) | 0.61 (1.66) | 0.18 (0.41) | 0.33 (1.05) |
| mir-210 | 0.41 (1.5) | 1.49 (3.46) | 0.14 (0.4) | 0.68 (2.23) |
| mir-214 | 0 (0) | 0 (0) | 0.01 (0.05) | 0 (0.03) |
| mir-218 | 0.03 (0.15) | 0 (0) | 0 (0) | 0.01 (0.09) |
| mir-224 | 0 (0) | 0 (0) | 0.04 (0.17) | 0.01 (0.1) |
| mir-23 | 0.42 (0.97) | 0.26 (0.6) | 0.22 (0.41) | 0.3 (0.69) |
| mir-24 | 0.21 (0.44) | 0.13 (0.3) | 0.1 (0.22) | 0.15 (0.33) |
| mir-26 | 33.95 (15.04) | 35.26 (13.11) | 32.99 (7.73) | 34.07 (12.18) |
| mir-28 | 8.2 (3.78) | 7.68 (4.77) | 10.76 (23.76) | 8.88 (13.99) |
| mir-29 | 5.66 (3.95) | 5.51 (2.56) | 4.93 (2.81) | 5.37 (3.12) |
| mir-290 | 0 (0) | 0 (0) | 0.03 (0.14) | 0.01 (0.08) |
| mir-296 | 0 (0) | 0.11 (0.49) | 0 (0) | 0.04 (0.28) |
| mir-3180 | 0.02 (0.11) | 0 (0) | 0.06 (0.28) | 0.03 (0.17) |
| mir-320 | 0.74 (1.15) | 0.47 (0.78) | 0.82 (1.34) | 0.68 (1.11) |
| mir-33 | 0.01 (0.06) | 0.04 (0.17) | 0.02 (0.1) | 0.02 (0.12) |
| mir-331 | 5.34 (5.31) | 4.07 (3.57) | 3.35 (3.53) | 4.25 (4.23) |
| mir-338 | 1.45 (1.66) | 1.55 (2.17) | 1.07 (1.21) | 1.36 (1.71) |
| mir-34 | 0.51 (1.77) | 0.49 (1.31) | 1.83 (3.75) | 0.94 (2.55) |
| mir-340 | 0.01 (0.05) | 0 (0) | 0 (0) | 0 (0.03) |
| mir-342 | 0 (0) | 0.02 (0.1) | 0 (0) | 0.01 (0.06) |
| mir-345 | 6.98 (7.32) | 4.14 (3.53) | 5.2 (5.86) | 5.44 (5.81) |
| mir-365 | 0.44 (1.5) | 0.33 (0.6) | 0.35 (0.95) | 0.37 (1.06) |
| mir-375 | 0.14 (0.32) | 0.18 (0.38) | 1.03 (3.51) | 0.45 (2.06) |
| mir-425 | 6.94 (2.85) | 5.84 (2.51) | 7 (4.82) | 6.59 (3.52) |
| mir-449 | 0.07 (0.29) | 0.04 (0.12) | 0.77 (2.29) | 0.29 (1.36) |
| mir-489 | 0 (0) | 0.03 (0.15) | 0 (0) | 0.01 (0.09) |
| mir-491 | 0 (0) | 0.13 (0.58) | 0 (0) | 0.04 (0.34) |
| mir-492 | 0 (0) | 0 (0) | 1.59 (7.1) | 0.53 (4.1) |
| mir-500 | 2.22 (2.69) | 2.12 (1.59) | 1.79 (1.9) | 2.04 (2.08) |
| mir-505 | 3.22 (2.83) | 2.68 (3.79) | 2.91 (3.43) | 2.94 (3.32) |
| mir-544 | 0.3 (1.34) | 0.08 (0.25) | 0.19 (0.68) | 0.19 (0.87) |
| mir-548 | 0.32 (0.69) | 0.31 (0.86) | 0.81 (2.26) | 0.48 (1.45) |
| mir-550 | 0.43 (1.1) | 0.61 (1.47) | 0.36 (0.84) | 0.46 (1.15) |
| mir-551 | 0.01 (0.05) | 0.05 (0.17) | 0 (0) | 0.02 (0.1) |
| mir-552 | 0.01 (0.06) | 0 (0) | 0 (0) | 0 (0.04) |
| mir-562 | 0.05 (0.16) | 0 (0) | 0.6 (2.68) | 0.22 (1.55) |
| mir-572 | 0 (0) | 0.05 (0.23) | 0 (0) | 0.02 (0.13) |
| mir-574 | 20.93 (14.57) | 15.29 (10.63) | 16.91 (15.11) | 17.71 (13.56) |
| mir-576 | 2.81 (5.95) | 1.47 (1.5) | 2.71 (4.5) | 2.33 (4.36) |
| mir-581 | 0 (0) | 0.01 (0.06) | 0.04 (0.17) | 0.02 (0.1) |
| mir-582 | 0.23 (0.45) | 0.8 (1.24) | 0.31 (0.41) | 0.45 (0.82) |
| mir-584 | 1.27 (1.63) | 2.77 (6.44) | 2.41 (2.84) | 2.15 (4.15) |
| mir-589 | 0.37 (0.73) | 0.37 (0.68) | 0.3 (0.52) | 0.35 (0.64) |
| mir-597 | 0.04 (0.12) | 0.08 (0.3) | 0.19 (0.6) | 0.1 (0.39) |
| mir-598 | 14.96 (15.14) | 7.66 (6.68) | 11.74 (10.09) | 11.46 (11.4) |
| mir-612 | 0.03 (0.13) | 0.02 (0.08) | 0 (0) | 0.02 (0.09) |
| mir-616 | 0.05 (0.17) | 0.2 (0.4) | 0.16 (0.41) | 0.14 (0.34) |
| mir-618 | 1.2 (1.61) | 1.32 (1.54) | 0.66 (0.95) | 1.06 (1.41) |
| mir-624 | 0.19 (0.39) | 0.52 (0.95) | 0.27 (0.5) | 0.33 (0.67) |
| mir-628 | 0.9 (1.29) | 2.33 (3.55) | 0.67 (0.77) | 1.3 (2.31) |
| mir-632 | 0 (0) | 0.05 (0.23) | 0.01 (0.06) | 0.02 (0.14) |
| mir-636 | 0 (0) | 0.08 (0.36) | 0.03 (0.15) | 0.04 (0.23) |
| mir-639 | 0.1 (0.46) | 0 (0) | 0 (0) | 0.03 (0.26) |
| mir-642 | 1.05 (1.88) | 1.27 (2.29) | 0.46 (0.61) | 0.93 (1.75) |
| mir-643 | 0.06 (0.25) | 0.1 (0.27) | 0.01 (0.06) | 0.06 (0.21) |
| mir-650 | 0 (0) | 0.23 (0.91) | 0 (0) | 0.08 (0.53) |
| mir-651 | 0.39 (1.08) | 0.39 (1.31) | 0.15 (0.33) | 0.31 (0.99) |
| mir-671 | 2.89 (3.29) | 1.47 (1.67) | 1.71 (3.6) | 2.02 (2.99) |
| mir-672 | 0 (0) | 0.04 (0.2) | 0 (0) | 0.01 (0.12) |
| mir-675 | 0.01 (0.06) | 0 (0) | 0 (0) | 0 (0.03) |
| mir-692 | 0.01 (0.06) | 0.16 (0.4) | 0.3 (1.34) | 0.16 (0.8) |
| mir-720 | 0 (0) | 0.02 (0.1) | 3.17 (14.19) | 1.07 (8.19) |
| mir-744 | 18.41 (20.9) | 13.73 (12.2) | 14.16 (16.82) | 15.43 (16.86) |
| mir-765 | 0.13 (0.46) | 0.09 (0.36) | 0.17 (0.33) | 0.13 (0.38) |
| mir-885 | 1.23 (1.96) | 1.15 (2.34) | 0.72 (0.98) | 1.03 (1.83) |
| mir-887 | 0.02 (0.11) | 0.02 (0.1) | 0.02 (0.08) | 0.02 (0.1) |
| mir-938 | 0 (0) | 0 (0) | 0.05 (0.19) | 0.02 (0.11) |
| mir-941 | 13.61 (10.91) | 7.37 (4.13) | 7.17 (4.16) | 9.38 (7.65) |
| mir-944 | 0.19 (0.45) | 0.6 (0.96) | 0.91 (1.27) | 0.57 (0.99) |
| mir-1846 | 4.96 (6.31) | 1.67 (5.15) | 11.6 (27.8) | 6.08 (16.96) |
| mir-811 | 18.1 (17.8) | 12.64 (23.67) | 39.74 (78.61) | 23.49 (49.11) |

* mean (standard deviation)

**Supplementary Table 7 | Spearman’s correlation coefficients between micro-RNA data and features of viral dynamics.**

| **Micro-RNA** | **Duration of viral shedding** | | **Peak viral load** | | **Up-slope** | | **Down-slope** | |
| --- | --- | --- | --- | --- | --- | --- | --- | --- |
|  | Spearman’s $\rho$ | p-value | Spearman’s $\rho$ | p-value | Spearman’s $\rho$ | p-value | Spearman’s $\rho$ | p-value |
| mir-10 | -0.16 | 0.9507 | -0.31 | 0.2323 | 0.06 | 0.944 | 0.07 | 0.9867 |
| mir-103 | 0.14 | 0.9507 | 0.06 | 0.8198 | 0.07 | 0.9321 | -0.16 | 0.9867 |
| mir-1180 | -0.08 | 0.9982 | 0.2 | 0.597 | -0.16 | 0.8616 | 0.12 | 0.9867 |
| mir-1226 | -0.03 | 0.9982 | 0.09 | 0.7441 | 0 | 0.993 | 0.03 | 0.9867 |
| mir-1253 | -0.16 | 0.9507 | 0.02 | 0.9244 | -0.17 | 0.8616 | 0.17 | 0.9867 |
| mir-1255 | -0.33 | 0.6991 | -0.09 | 0.7441 | -0.08 | 0.9247 | 0.34 | 0.7121 |
| mir-128 | 0.04 | 0.9982 | 0.12 | 0.6894 | -0.09 | 0.9247 | -0.01 | 0.9867 |
| mir-1296 | -0.01 | 0.9982 | 0.17 | 0.6225 | 0.02 | 0.9497 | 0.02 | 0.9867 |
| mir-130 | 0.13 | 0.9507 | 0.18 | 0.597 | 0.02 | 0.9497 | -0.07 | 0.9867 |
| mir-145 | 0.12 | 0.9553 | 0.01 | 0.9822 | 0.08 | 0.9247 | -0.12 | 0.9867 |
| mir-146 | 0.17 | 0.9507 | 0.16 | 0.635 | 0.09 | 0.9247 | -0.15 | 0.9867 |
| mir-148 | -0.02 | 0.9982 | 0.25 | 0.4697 | -0.11 | 0.9029 | 0.04 | 0.9867 |
| mir-154 | -0.03 | 0.9982 | 0.14 | 0.635 | -0.18 | 0.8616 | 0.07 | 0.9867 |
| mir-16 | 0.25 | 0.7309 | 0.07 | 0.7731 | 0.18 | 0.8616 | -0.27 | 0.9509 |
| mir-17 | 0.02 | 0.9982 | 0.04 | 0.87 | -0.04 | 0.9497 | -0.04 | 0.9867 |
| mir-181 | 0.08 | 0.9982 | 0.06 | 0.8198 | 0 | 0.9925 | -0.07 | 0.9867 |
| mir-185 | -0.07 | 0.9982 | -0.19 | 0.597 | 0.08 | 0.9247 | 0 | 0.9918 |
| mir-19 | -0.09 | 0.9982 | 0.25 | 0.4697 | -0.16 | 0.8616 | 0.16 | 0.9867 |
| mir-191 | -0.01 | 0.9982 | -0.14 | 0.635 | 0.12 | 0.9029 | -0.04 | 0.9867 |
| mir-192 | -0.03 | 0.9982 | -0.06 | 0.8198 | 0.19 | 0.8616 | 0.01 | 0.9867 |
| mir-193 | -0.17 | 0.9507 | -0.1 | 0.7441 | 0.02 | 0.9497 | 0.16 | 0.9867 |
| mir-204 | -0.02 | 0.9982 | 0.28 | 0.3167 | -0.25 | 0.8616 | 0.06 | 0.9867 |
| mir-210 | -0.04 | 0.9982 | 0.21 | 0.597 | -0.21 | 0.8616 | 0.06 | 0.9867 |
| mir-214 | 0.14 | 0.9507 | -0.02 | 0.9244 | 0.14 | 0.9029 | -0.13 | 0.9867 |
| mir-218 | -0.16 | 0.9507 | 0.02 | 0.9244 | -0.17 | 0.8616 | 0.17 | 0.9867 |
| mir-224 | 0.22 | 0.8213 | 0.16 | 0.635 | 0.22 | 0.8616 | -0.2 | 0.9867 |
| mir-23 | -0.02 | 0.9982 | 0.08 | 0.7719 | 0.11 | 0.9029 | 0.03 | 0.9867 |
| mir-24 | -0.1 | 0.9982 | -0.08 | 0.7731 | 0.05 | 0.944 | 0.11 | 0.9867 |
| mir-26 | 0.07 | 0.9982 | -0.06 | 0.8198 | 0.01 | 0.9736 | -0.09 | 0.9867 |
| mir-28 | -0.27 | 0.6991 | -0.11 | 0.6894 | -0.09 | 0.9247 | 0.23 | 0.9867 |
| mir-29 | -0.07 | 0.9982 | 0 | 0.9926 | -0.12 | 0.9029 | 0.06 | 0.9867 |
| mir-290 | 0.2 | 0.9175 | 0.11 | 0.6894 | 0.19 | 0.8616 | -0.17 | 0.9867 |
| mir-296 | -0.02 | 0.9982 | 0.08 | 0.7731 | -0.15 | 0.8892 | 0.04 | 0.9867 |
| mir-3180 | -0.02 | 0.9982 | -0.2 | 0.597 | 0.12 | 0.9029 | -0.01 | 0.9867 |
| mir-320 | -0.01 | 0.9982 | -0.12 | 0.6615 | -0.03 | 0.9497 | 0 | 0.9979 |
| mir-33 | 0 | 0.9982 | -0.05 | 0.8198 | 0.13 | 0.9029 | 0.01 | 0.9867 |
| mir-331 | -0.03 | 0.9982 | -0.05 | 0.8373 | -0.05 | 0.944 | -0.01 | 0.9867 |
| mir-338 | 0 | 0.9982 | 0.16 | 0.635 | 0.08 | 0.9247 | 0.02 | 0.9867 |
| mir-34 | 0.27 | 0.6991 | 0.12 | 0.6615 | 0.12 | 0.9029 | -0.24 | 0.9867 |
| mir-340 | -0.13 | 0.9507 | -0.11 | 0.6894 | 0.05 | 0.944 | 0.11 | 0.9867 |
| mir-342 | 0.01 | 0.9982 | 0.2 | 0.597 | -0.18 | 0.8616 | 0.01 | 0.9867 |
| mir-345 | -0.15 | 0.9507 | -0.13 | 0.6517 | -0.08 | 0.9247 | 0.11 | 0.9867 |
| mir-365 | 0.06 | 0.9982 | 0 | 0.9926 | 0.09 | 0.9247 | -0.09 | 0.9867 |
| mir-375 | 0.13 | 0.9507 | 0.05 | 0.8198 | 0.07 | 0.9321 | -0.09 | 0.9867 |
| mir-425 | -0.11 | 0.985 | -0.14 | 0.635 | 0.01 | 0.9724 | 0.06 | 0.9867 |
| mir-449 | 0.29 | 0.6991 | 0.14 | 0.635 | 0.17 | 0.8616 | -0.27 | 0.9509 |
| mir-489 | -0.11 | 0.985 | 0.09 | 0.7441 | -0.2 | 0.8616 | 0.14 | 0.9867 |
| mir-491 | 0.07 | 0.9982 | 0.14 | 0.635 | -0.08 | 0.9247 | -0.07 | 0.9867 |
| mir-492 | 0.16 | 0.9507 | -0.01 | 0.9867 | 0.17 | 0.8616 | -0.14 | 0.9867 |
| mir-500 | -0.03 | 0.9982 | 0.16 | 0.635 | -0.08 | 0.9247 | 0.07 | 0.9867 |
| mir-505 | 0 | 0.9982 | -0.22 | 0.597 | 0.16 | 0.8616 | -0.07 | 0.9867 |
| mir-544 | 0.11 | 0.985 | -0.05 | 0.8198 | -0.02 | 0.9497 | -0.13 | 0.9867 |
| mir-548 | 0.03 | 0.9982 | -0.11 | 0.6894 | 0.02 | 0.9497 | -0.04 | 0.9867 |
| mir-550 | -0.1 | 0.9982 | -0.02 | 0.9244 | -0.03 | 0.9497 | 0.14 | 0.9867 |
| mir-551 | -0.05 | 0.9982 | 0.17 | 0.6225 | -0.17 | 0.8616 | 0.06 | 0.9867 |
| mir-552 | -0.09 | 0.9982 | -0.07 | 0.8012 | -0.02 | 0.9497 | 0.09 | 0.9867 |
| mir-562 | -0.09 | 0.9982 | -0.17 | 0.6207 | -0.08 | 0.9247 | 0.06 | 0.9867 |
| mir-572 | 0.04 | 0.9982 | 0.13 | 0.6605 | -0.1 | 0.9247 | -0.02 | 0.9867 |
| mir-574 | -0.14 | 0.9507 | -0.18 | 0.597 | -0.02 | 0.9497 | 0.08 | 0.9867 |
| mir-576 | 0 | 0.9982 | 0.09 | 0.7511 | -0.07 | 0.9331 | 0.02 | 0.9867 |
| mir-581 | 0.17 | 0.9507 | 0.14 | 0.635 | 0.13 | 0.9029 | -0.17 | 0.9867 |
| mir-582 | 0.07 | 0.9982 | 0.2 | 0.597 | -0.17 | 0.8616 | -0.01 | 0.9867 |
| mir-584 | 0.15 | 0.9507 | 0.04 | 0.8503 | -0.07 | 0.9331 | -0.16 | 0.9867 |
| mir-589 | 0.06 | 0.9982 | 0.13 | 0.6517 | 0.06 | 0.944 | -0.03 | 0.9867 |
| mir-597 | -0.03 | 0.9982 | -0.1 | 0.724 | 0.05 | 0.944 | 0.03 | 0.9867 |
| mir-598 | -0.08 | 0.9982 | -0.32 | 0.2323 | 0.11 | 0.9029 | 0 | 0.9918 |
| mir-612 | -0.07 | 0.9982 | 0.1 | 0.724 | -0.17 | 0.8616 | 0.1 | 0.9867 |
| mir-616 | 0.13 | 0.9507 | 0.19 | 0.597 | -0.07 | 0.936 | -0.08 | 0.9867 |
| mir-618 | -0.13 | 0.9507 | 0.02 | 0.9353 | -0.05 | 0.944 | 0.13 | 0.9867 |
| mir-624 | 0.03 | 0.9982 | 0.18 | 0.597 | -0.13 | 0.9029 | 0.01 | 0.9867 |
| mir-628 | 0.03 | 0.9982 | 0.18 | 0.597 | -0.08 | 0.9247 | 0.01 | 0.9867 |
| mir-632 | 0.08 | 0.9982 | 0.29 | 0.3167 | -0.13 | 0.9029 | -0.02 | 0.9867 |
| mir-636 | 0.15 | 0.9507 | 0.09 | 0.7511 | 0.04 | 0.9497 | -0.14 | 0.9867 |
| mir-639 | -0.06 | 0.9982 | -0.19 | 0.597 | 0.2 | 0.8616 | -0.01 | 0.9867 |
| mir-642 | -0.02 | 0.9982 | 0.09 | 0.7511 | 0.05 | 0.944 | 0.03 | 0.9867 |
| mir-643 | -0.02 | 0.9982 | 0.24 | 0.4955 | -0.17 | 0.8616 | 0.05 | 0.9867 |
| mir-650 | -0.01 | 0.9982 | 0.07 | 0.8012 | -0.12 | 0.9029 | 0.03 | 0.9867 |
| mir-651 | -0.01 | 0.9982 | 0.09 | 0.7511 | -0.04 | 0.9497 | 0.05 | 0.9867 |
| mir-671 | -0.13 | 0.9507 | -0.05 | 0.8198 | 0.04 | 0.944 | 0.1 | 0.9867 |
| mir-672 | 0.01 | 0.9982 | 0.2 | 0.597 | -0.18 | 0.8616 | 0.01 | 0.9867 |
| mir-675 | -0.22 | 0.8213 | -0.14 | 0.635 | 0.1 | 0.9247 | 0.22 | 0.9867 |
| mir-692 | -0.03 | 0.9982 | 0.05 | 0.8198 | -0.2 | 0.8616 | 0.05 | 0.9867 |
| mir-720 | 0.12 | 0.9553 | 0.14 | 0.635 | 0 | 0.993 | -0.1 | 0.9867 |
| mir-744 | -0.05 | 0.9982 | -0.2 | 0.597 | 0.06 | 0.944 | -0.03 | 0.9867 |
| mir-765 | 0.25 | 0.7309 | -0.08 | 0.7559 | 0.38 | 0.2349 | -0.26 | 0.9509 |
| mir-885 | -0.07 | 0.9982 | 0 | 0.9926 | 0.01 | 0.9724 | 0.09 | 0.9867 |
| mir-887 | 0.09 | 0.9982 | 0.14 | 0.635 | 0.03 | 0.9497 | -0.06 | 0.9867 |
| mir-938 | 0.22 | 0.8213 | 0.14 | 0.635 | 0.14 | 0.9029 | -0.17 | 0.9867 |
| mir-941 | -0.22 | 0.8213 | -0.37 | 0.1235 | 0.12 | 0.9029 | 0.08 | 0.9867 |
| mir-944 | 0.28 | 0.6991 | 0.37 | 0.1235 | -0.01 | 0.9925 | -0.2 | 0.9867 |
| mir-1846 | -0.17 | 0.9507 | -0.53 | 0.0011 | 0.2 | 0.8616 | 0.09 | 0.9867 |
| mir-811 | -0.13 | 0.9507 | -0.31 | 0.2323 | 0.05 | 0.944 | 0.06 | 0.9867 |

**Reference**

1	Ke, R. *et al.* Daily longitudinal sampling of SARS-CoV-2 infection reveals substantial heterogeneity in infectiousness. *Nat Microbiol* **7**, 640-652 (2022). <https://doi.org:10.1038/s41564-022-01105-z>
